# Circadian Regulation of Vertebrate Cone Photoreceptor Function

**DOI:** 10.1101/2021.05.05.442714

**Authors:** Jingjing Zang, Matthias Gesemann, Jennifer Keim, Marijana Samardzija, Christian Grimm, Stephan C.F. Neuhauss

## Abstract

Eukaryotes generally display a circadian rhythm as an adaption to the reoccurring day/night cycle. This is particularly true for visual physiology that is directly affected by changing light conditions. Here we investigate the influence of the circadian rhythm on the expression and function of visual transduction cascade regulators in diurnal zebrafish and nocturnal mice. We focused on regulators of shut-off kinetics such as *recoverins, arrestins, opsin kinases*, and *GTPase-accelerating protein* that have direct effects on temporal vision. Transcript as well as protein levels of most analyzed genes show a robust circadian rhythm dependent regulation, which correlates with changes in photoresponse kinetics. Electroretinography demonstrates that photoresponse recovery in zebrafish is delayed in the evening and accelerated in the morning. This physiological rhythmicity is mirrored in visual behaviors, such as optokinetic and optomotor responses. Functional rhythmicity persists in continuous darkness, it is reversed by an inverted light cycle and disrupted by constant light. This is in line with our finding that orthologous gene transcripts from diurnal zebrafish and nocturnal mice are often expressed in an anti-phasic daily rhythm.

## Introduction

Circadian rhythms serve as endogenous clocks that molecularly support the daily occurring oscillations of physiology and ensuing behavior (Brown *et al*, 2019; Cahill, 2002; Frøland Steindal & Whitmore, 2019; Golombek *et al*, 2014; Idda *et al*, 2012; Ukai & Ueda, 2010; Vatine *et al*, 2011). It has long been recognized that the central pacemaker of circadian rhythms resides in dedicated brain regions, either the suprachiasmatic nucleus in mammals or the pineal gland in non-mammalian vertebrates. The rhythm is entrained by external stimuli (e.g. light) that directly acts on the core circadian transcriptional feedback loop. Multiple studies have shown that autonomous circadian clocks also exist in other brain regions and in peripheral tissues (Frøland Steindal & Whitmore, 2019; Idda *et al*., 2012; Vatine *et al*., 2011). This is particularly true for the retina, which generates its own circadian rhythm (Ko, 2020). In zebrafish this rhythmicity is reflected in a number of circadian adaptations, such as a higher response threshold in the morning (Li & Dowling, 1998), photoreceptor retinomotor movement in constant darkness (Menger *et al*, 2005) and cone photoreceptor synaptic ribbon disassembly at night (Emran *et al*, 2010). Such adaptations are also found in other animals such as mice where stronger electrical retinal coupling during the night (Jin *et al*, 2015; Li *et al*, 2009; Ribelayga *et al*, 2008), as well as slower dark adaptation of rods during the day was observed (Xue *et al*, 2015). The molecular mechanisms underlying these circadian dependent retinal regulations are still largely unknown.

In the vertebrate retina there are two different types of photoreceptors, namely rods and cones (Burns & Baylor, 2001; Fu & Yau, 2007). Rods function mainly during dim light conditions, whereas cones are characterized by lower sensitivity but faster response kinetics being important for daylight and color vision. About 92% of larval and 60% of adult photoreceptors in the zebrafish retina are cones (Allison *et al*, 2010; Fadool, 2003; Zimmermann *et al*, 2018). Although rods and cones generally use the same visual transduction cascade components, the individual reactions are typically mediated by photoreceptor type specific proteins.

Visual transduction commences by an opsin chromophore mediated absorption of photons, which triggers the activation of a second messenger cascade including the trimeric G-protein transducin. Activated transducin stimulates the effector enzyme phosphodiesterase (PDE), which leads to a reduction of intracellular cGMP levels, subsequently leading to the closure of CNG-gated cation channels resulting in a membrane potential change (Fain *et al*, 2001; Lamb & Pugh, 2006).

High-temporal resolution requires a tightly regulated termination of visual transduction (Chen *et al*, 2012; Matthews & Sampath, 2010; Zang & Matthews, 2012). This depends on the highly effective quenching of both the activated visual pigment (R*) and PDE-transducin complex (PDE*). R* are phosphorylated by a G-protein receptor kinase (GRK) before being completely deactivated by binding to arrestin. While GRK activity itself is controlled by recoverin (RCV) in a Ca^2+^-dependent manner (Zang & Neuhauss, 2018), the quenching of PDE* depends on the GTPase activity of its γ-subunit that is regulated by activator protein RGS9 (Regulator of G-protein Signaling 9)(Krispel *et al*, 2006).

We now show that the expression levels of these important regulators of cone visual transduction decay are modulated by the circadian clock. Moreover, these periodic fluctuations are reflected in oscillating protein levels that correlate with the rhythmicity in visual physiology and behavior observed in zebrafish. Interestingly, we found that the expression of a selection of mouse orthologes of the investigated regulatory genes are also modulated by the circadian clock. However, the periodicity was opposite to zebrafish, fitting the nocturnal lifestyle of mice.

## Results

### Expression levels of key genes involved in shaping visual transduction decay are regulated by the circadian clock

In order to determine the influence of the circadian clock on visual behavior, we analyzed gene expression levels of key visual transduction regulators over a 24-hour-period using quantitative real time PCR (qRT-PCR). Eyes from larval (5 dpf) and adult zebrafish that were kept under a normal light cycle (LD), as well as eyes from 5 dpf larvae kept in continuous darkness (DD), were collected every 3 hours over a period of 24 hours and subsequently analyzed. Apart from *rcv2a* which seems to have no fluctuating transcript levels in larvae (Figure 1I), expression levels of the other *recoverins* (*rcv1a, rcv1b* which is absent from larval retina and *rcv2b*), *G-protein receptor kinases* (*grk7a* and *grk7b*), *arrestins* (*arr3a* and *arr3b*) and *regulator of G-protein signaling 9* (*rgs9a*) were clearly oscillating (Figure 1, supplement table 1). In many cases, transcripts were most abundant in the morning (*grk7a, grk7b* and *rgs9a*) or at midday (*rcv2b, arr3a* and *arr3b*), subsequently declined throughout the day and recovering during the night. For instance, in adult zebrafish eyes *grk7a* expression levels decreased by around 98% from peak expression to lowest expression levels (Figure 1A). *In situ* hybridization (ISH) analysis using digoxigenin labeled RNA probes validated our qRT-PCR results (Figure 1-figure supplement 2&3).

**Table 1:**
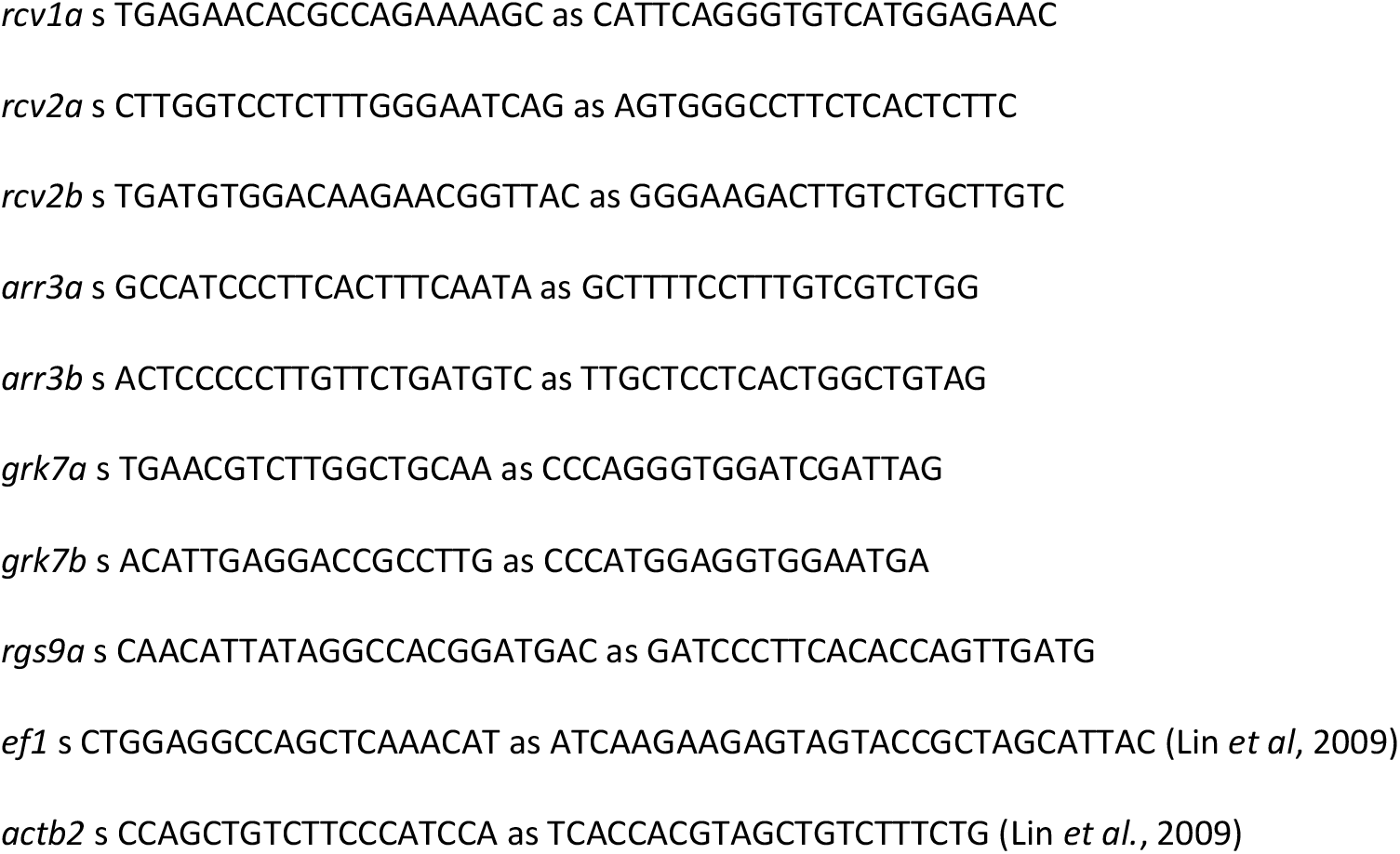
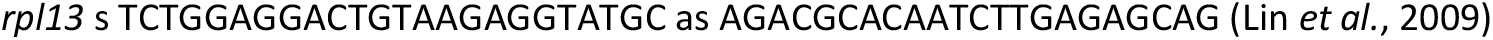
Sequences of primers used for q-RT PCR.

**Table 2.**
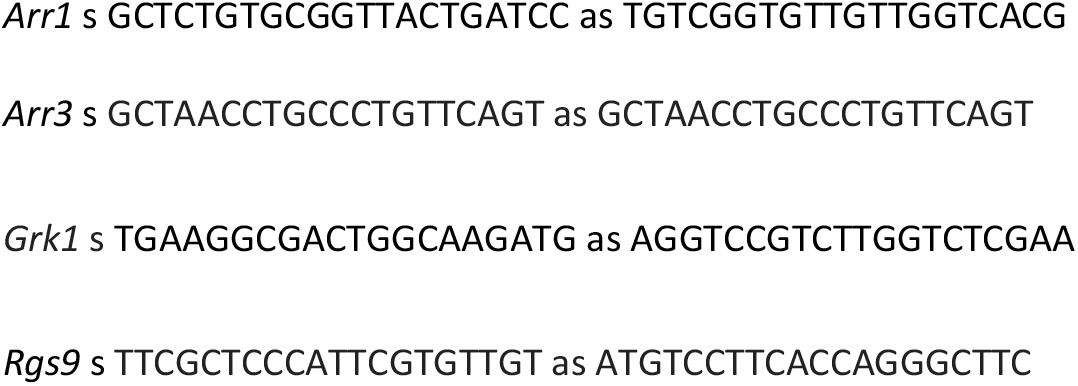
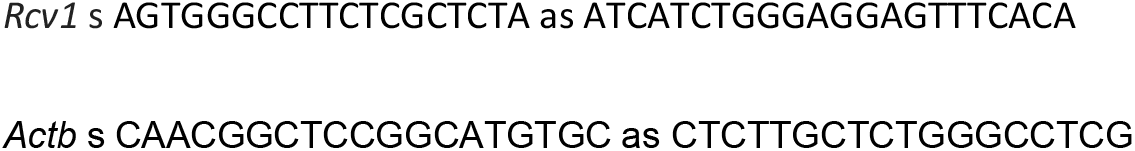
Mouse Primer Sequences.

**Figure 1.**
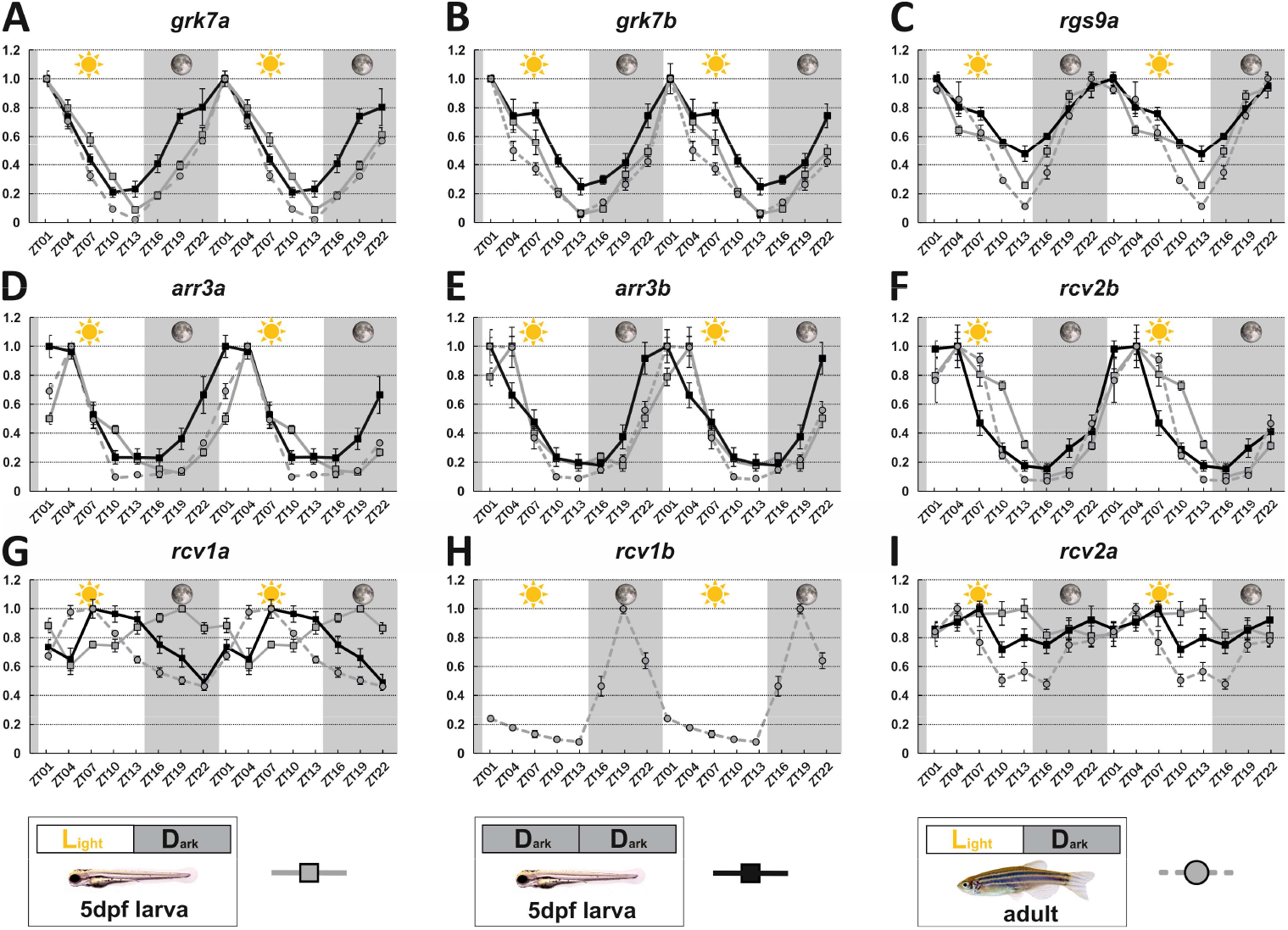
Key visual transduction decay gene transcripts are under circadian control. mRNA levels of visual transduction decay genes in the eye of adult and larval zebrafish were measured by qRT-PCR over a 24-hour-period and the data was duplicated in a 48-hour-time scale. (A-I). Eye tissues from larval fish either raised under a normal light/dark cycle (LD / gray squares) or in continuous darkness (DD / black squares) and from adult LD zebrafish (gray circles) were collected at eight different time points throughout the day. The name of the analyzed gene transcripts is given on top of each graph. The time point of collection is indicated along the x-axis with ZT01 being the time point one hour after the light was turned on. Dark periods are indicated by the moon symbol and highlighted in gray, whereas the periods under regular light conditions are indicated by the sun symbol and shown in white. For better orientation the different conditions are summarized at the bottom of the figure. Data represents the mean ± standard error of the mean (s.e.m) of three or more independent measurements. Statistics information is provided in Supplementary file 1.

Interestingly, two genes, namely *rcv1a* and *rcv2a*, displayed different expression profiles in larval and adult eyes (Figure 1G,I). While larval *rcv1a* mRNA transcript levels peaked around midnight, larval *rcv2a* transcript expression was non-cyclic (Figure 1G,I). However, this is in contrast to adult retinas where *rcv1a* and *rcv2a* transcripts were highest at midday (Figure 1G). An anti-phasic expression profile between larval and adult stages can also be observed for rod *arrestin* (*arras*) (Figure 1-figure supplement 4).

In order to establish that the daily expression changes of these transcripts are indeed regulated by the intrinsic circadian clock, we repeated our experiments in larvae kept in complete darkness (DD), eliminating light as an external factor. Under normal LD as well as DD conditions we obtained largely comparable results (Figure 1), although there was a 3-hour-phase shift in both *arr3a* and *arr3b* (Figure 1C&D).

### Corresponding retinal genes in nocturnal mice display an anti-phasic expression pattern

As zebrafish are diurnal animals having a cone dominant retina, we wondered if the observed circadian regulation of visual transduction gene transcripts is also seen in the rod dominant retina of nocturnal mice. We selected mouse *Grk1*, the only visual grk gene in mice (Chen *et al*, 1999; Wada *et al*, 2006), the sole recoverin (Chen *et al*., 2012) and Rgs9 (Krispel *et al*., 2006) genes and the two arrestins *Arr1* and *Arr3*, as the counterparts for the above mentioned zebrafish genes for our analysis.

Expression of all five regulators fluctuated in a 24-hour-period (Figure 2), being highest at the beginning of the dark period (ZT13) for the two *arrestins* (Figure. 2A&B), or around midnight (ZT17) for *Grk1, Rgs1* and *Rcv1*. All of them displayed minimal transcript levels early during the day. This oscillation pattern shows a clear anti-phasic relationship with the cyclic fluctuation of the corresponding zebrafish transcripts. Curiously, the amplitude of gene fluctuation in adult zebrafish retina was generally larger than in the mouse retina (Figures 1&2).

**Figure 2.**
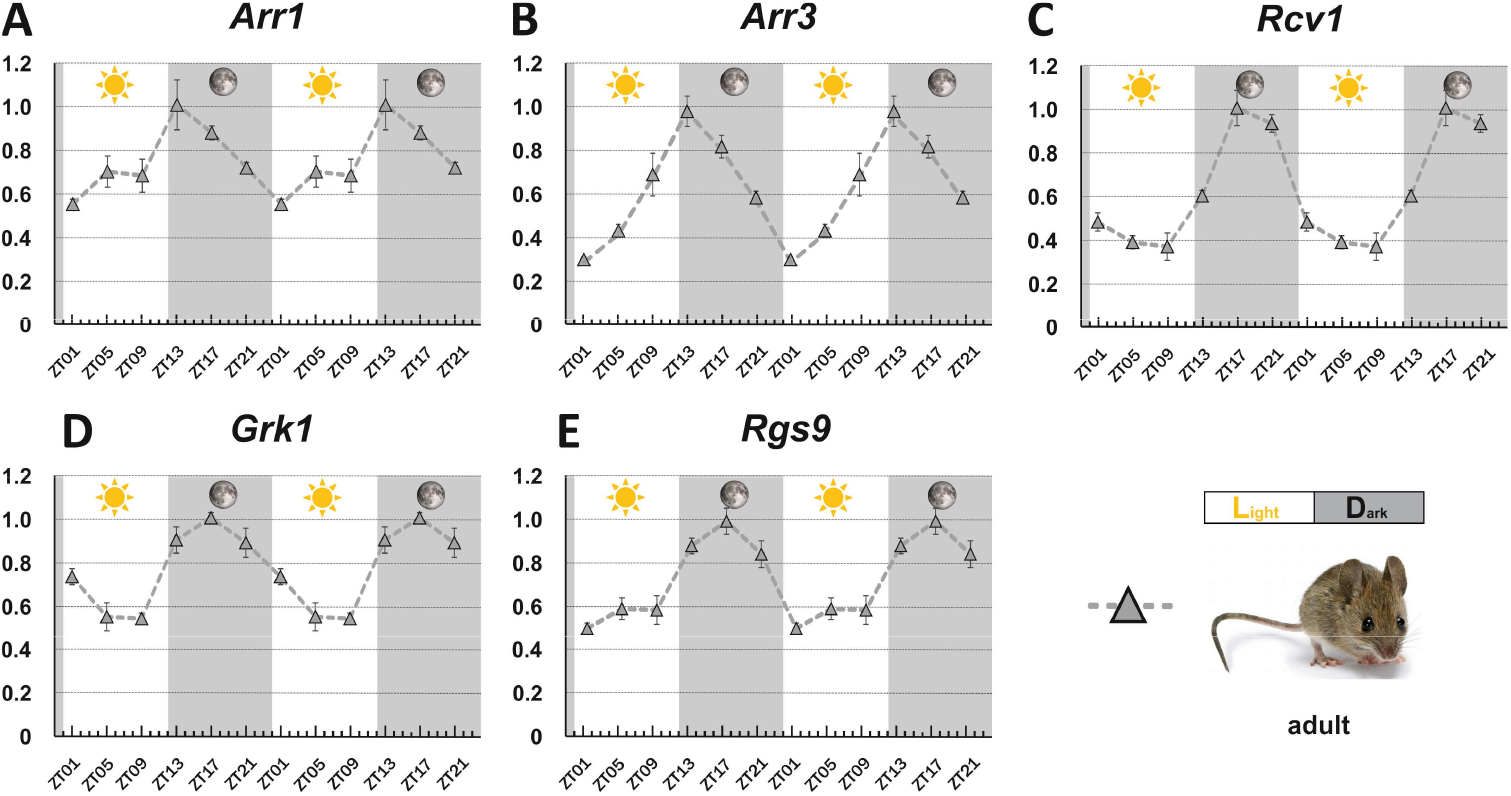
Circadian regulation of key visual transduction genes in nocturnal mice is reversed. Transcript levels of indicated mouse genes were measured using qRT-PCR on retinal tissue of 12-week-old wildtype mice. The time point of collection is indicated along the x-axis with ZT01 being the time point one hour after the light was turned on. The data was duplicated in a 48-hour-time scale. Dark periods are indicated by the moon symbol and highlighted in gray, whereas the periods under regular light conditions are indicated by the sun symbol and shown in white. Data represents the mean ± s.e.m of three independent measurements. One-way ANOVA was performed by GraphPad Prism 8. p=0.018 at ZT13, p=0.008 at ZT17 and p=0.001 at ZT21 compared with the lowest point at ZT1 (A). p=0.015 at ZT5, p=0.015 at ZT9, p<0.001 at ZT13, p<0.001 at ZT17 and p<0.001 at ZT21 compared with the lowest point at ZT1 (B). p=0.014 at ZT13, p=0.022 at ZT17 and p<0.001 at ZT21 compared with the lowest level at ZT9 (C). p=0.011 at ZT1, p=0.005 at ZT13, p<0.001 at ZT17 and p=0.008 at ZT21 compared with the lowest point at ZT9 (D). p<0.001 at ZT13, p=0.001 at ZT17 and p=0.007 at ZT21compared with the lowest level atZT1 (E).

Despite the rhythmic expression in both zebrafish and mice, our bioinformatics search did not discover conserved transcription factor binding sites of known core circadian regulators in any of the tested genes (Figure 2-figure supplement 1).

### Levels of key visual transduction regulator proteins fluctuate in the zebrafish retina

While mRNA half-life is typically in the range of minutes, protein turnover rates can range from minutes to days, explaining why fluctuation of mRNA levels are not always reflected in time-shifted oscillations at the protein level (Cunningham & Gonzalez-Fernandez, 2000; Stenkamp *et al*, 2005). However, as regulatory proteins often have turnover rates of only a few hours we were examining whether RNA oscillations are mirrored by corresponding protein level fluctuations. In order to assess protein levels, we generated paralogue-specific antibodies against GRK7a and ARR3a. Quantitative Western blot analysis indicated periodic changes in protein levels for both proteins. Peak expression was shifted 6 to 12 hours between RNA and protein levels (Figure 3A, 3B). ARR3a reached its highest and lowest levels at ZT7 and ZT22, respectively, whereas GRK7a maintained relatively high levels throughout the day having the lowest concentrations around midnight. Hence, mRNA circadian oscillations in the zebrafish retina are largely conserved at the protein level with a time shift.

**Figure 3.**
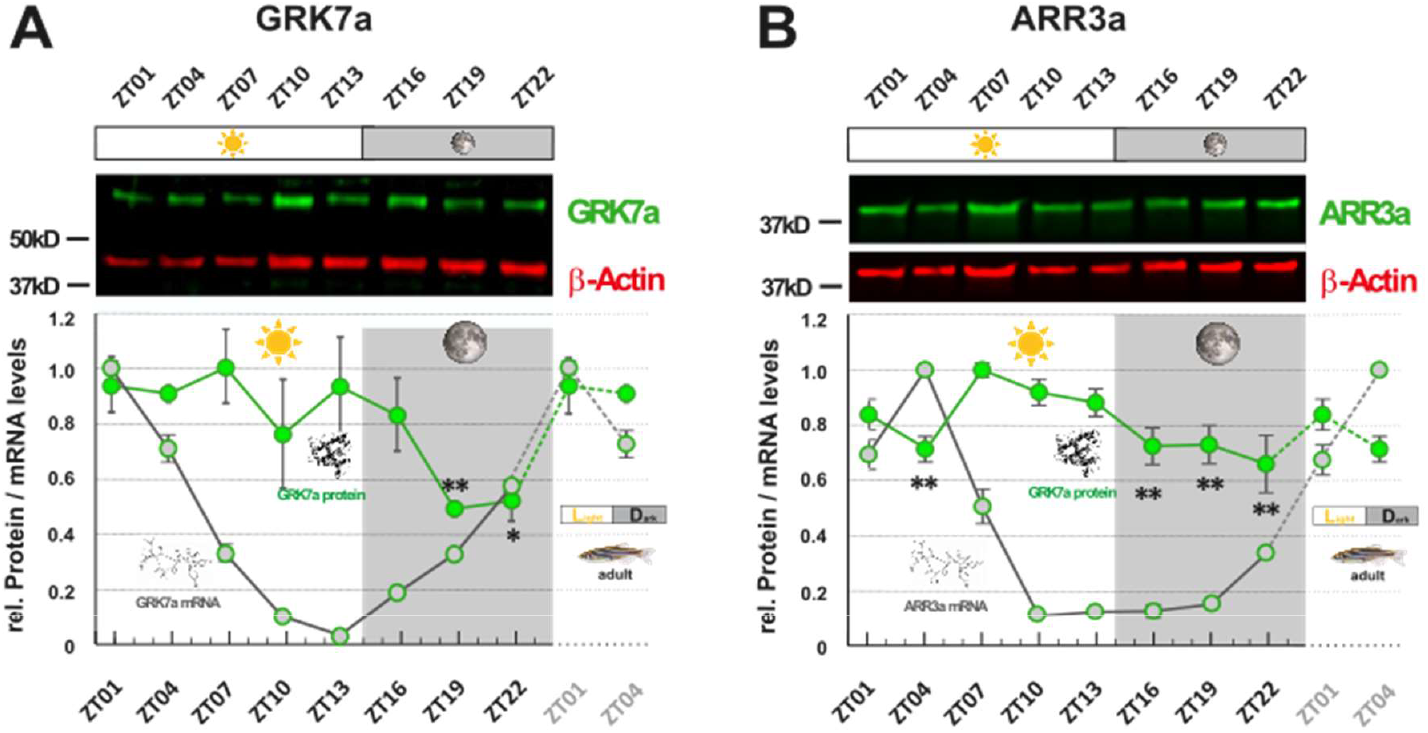
GRK7a and ARR3a protein levels show daily changes in adult zebrafish eyes. GRK7a (A) and ARR3a (B) protein levels were quantified using Western blot analysis. β-Actin was used as a loading control. While mRNA transcript levels (gray circles / RNA structure) were lowest in the evening (ZT10 and ZT13, respectively), lowest protein expression levels (green circles / protein structure) were tailing RNA expression levels by around 6 to 12 hours, reaching lowest levels in the middle of the night at around ZT19. The time point of collection is indicated along the x-axis with ZT01 being the time point one hour after the light was turned on. Dark periods are indicated by the moon symbol and highlighted in gray, whereas the periods under regular light conditions are indicated by the sun symbol and shown in white. One-way ANOVA was performed by SPSS (IBM, version 26.0). p=0.009 at ZT19 and p=0.013 at ZT22 compared with the highest level at ZT7 in (A). p= 0.003 at ZT04, p= 0.004 at ZT16, p= 0.005 at ZT19 and p= 0.001 at ZT22 compared with the highest level at ZT7 in (B). Data represents the mean ± s.e.m of three independent measurements in (A) and four independent measurements in (B). * p<0.05 **p<0.01

### Larval cone response recovery is delayed in the evening

We next asked whether the observed protein and RNA level fluctuations have an impact on functional aspects of visual transduction. In the electroretinogram (ERG) the a-wave directly represents photoreceptor responses. Since in the zebrafish ERG it is largely masked by the larger b-wave, reflecting the depolarization of ON-bipolar cells, we used the b-wave amplitude as an indirect measure of the cone photoresponse (Figure 4A1). The protein products of the genes analyzed in our study are known to affect photoresponse recovery in zebrafish (Renninger *et al*, 2011; Rinner *et al*, 2005; Zang *et al*, 2015). Therefore, we assessed their function by using the ERG double flash paradigm. In this experimental set-up, the retina receives a conditioning flash followed by a probing flash of the same light intensity (Figure 4A1). The b-wave amplitude ratio of probing to conditioning response in relation to the interstimulus interval is a normalized read-out for the visual transduction recovery time (Figure 4A2). Photoreceptor recovery is complete when the two flashes evoke responses of equal amplitudes.

**Figure 4.**
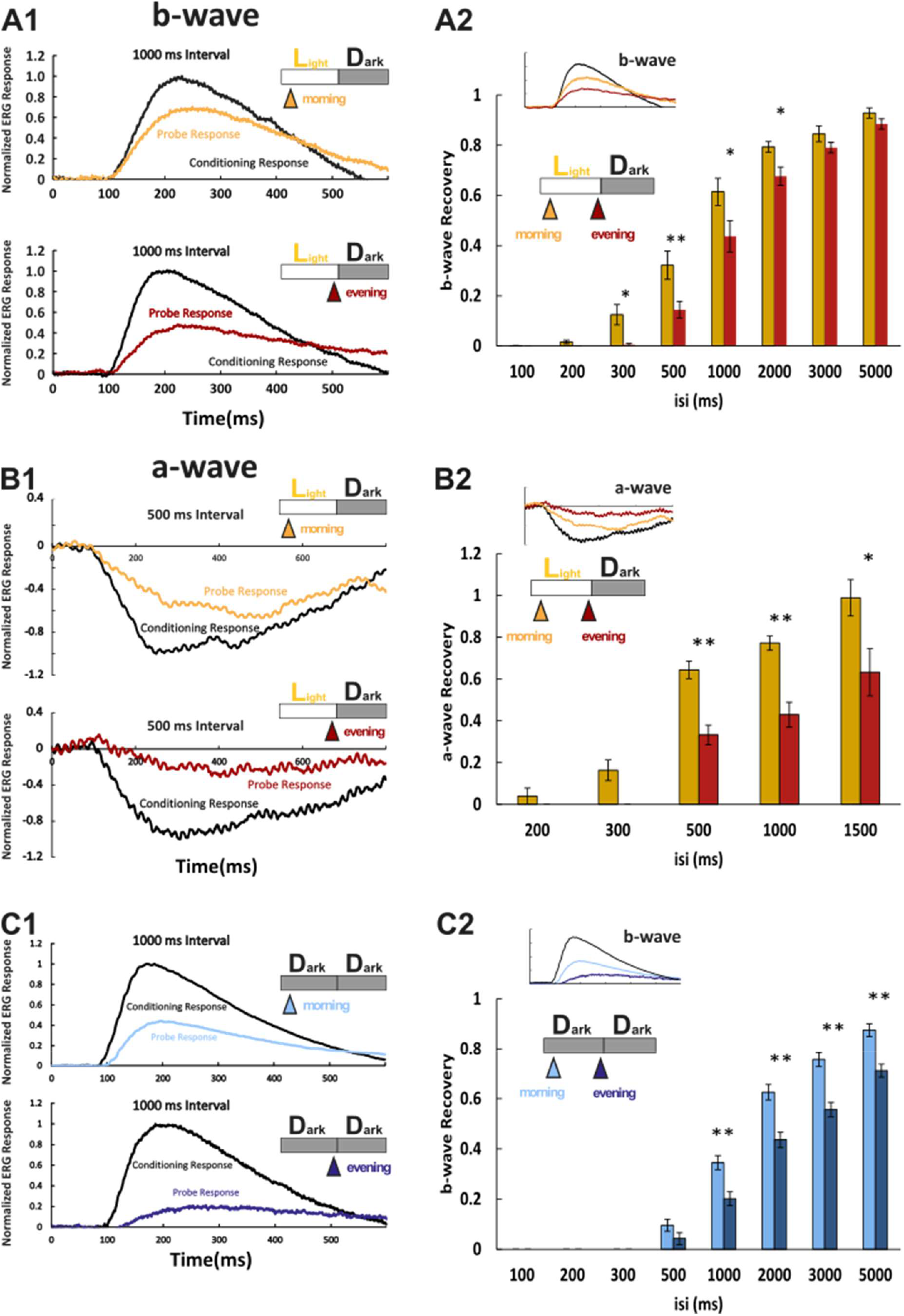
Larval cone photoresponse recovery is accelerated in the morning. **(A1)** Examples of LD larval ERG b-wave recordings. A conditioning flash (black line) was followed by a probing flash (yellow and red lines) that were separated by 1000 ms. While the yellow triangle and curve mark the probe response in the morning, the red triangle and curve represents the probe response recorded in the evening. Note that the probe response in the evening is clearly diminished. **(A2)** b-wave recovery as a function of the interstimulus interval (isi). At 500 ms up to 3000 ms isi b-wave recovery in the morning (yellow bars) is significantly enhanced when compared to corresponding recordings in the evening (red bars). Note that below 500 ms isi no b-wave recovery can be observed and that at an interval of 5 s complete recovery can also be found in the evening. Data are presented as mean ± s.e.m (n=18 in the morning; n=14 in the evening) of three independent experiments. Student’s t-test was used to compare the response in the morning and in the evening. p=0.011 at 300 ms isi. p=0.010 at 500 ms isi. p=0.026 at 1000 ms isi. p=0.012 at 2000ms isi. * p<0.05 **p<0.01 **(B1)** Examples of LD larval ERG a-wave recordings under DL-TBOA and L-AP4 inhibition. Under b-wave blocking conditions a conditioning flash (black line) is followed by a probing flash (yellow and red lines) that were separated by 500 ms. The yellow triangle and curve mark the probe response in the morning, whereas the red triangle and curve represents the probe response recorded in the evening. Note that also the a-wave response recovery is significantly reduced in the evening. **(B2)** a-wave recovery as a function of isi. At 300 ms up to 1500 ms isi a-wave recovery in the morning (yellow bars) is significantly enhanced when compared to corresponding recordings in the evening (red bars). Data are presented as mean ± s.e.m (n=11 in the morning; n=5 in the evening) of three independent experiments. Student’s t-test was used. p=0.003 at 500 ms isi. p=0.0003 at 1000ms isi. p=0.038 at 1500 ms isi. * p<0.05 **p<0.01 **(C1)** Examples of ERG b-wave recordings from a larva kept under constant darkness (DD). A conditioning flash (black line) was followed by a probing flash (light and dark blue lines) that were separated by 1000 ms. The light blue triangle and curve mark the probe response at the subjective morning, whereas the dark blue triangle and curve represents the probe response recorded at the subjective evening. **(C2)** b-wave recovery as a function of the isi is shown for larvae raised in continuous darkness DD. Even under continuous darkness visual function remains under circadian control as at 500 ms up to 3000 ms isi, the b-wave recovery in the subjective morning (light blue bars) is significantly enhanced when compared to corresponding recordings in the subjective evening (dark blue bars). Data are presented as mean ± s.e.m (n=17 in the morning; n=12 in the evening) of three independent experiments. Student’s t-test was used. p=0.0007 at 1000 ms isi. p= 0.002 at 2000 ms isi. p=0.0004 at 3000 ms isi. p=0.0006 at 5000 ms isi. * p<0.05 **p<0.01

Response recovery was significantly delayed in the evening in comparison to the morning (Figure 4A2). However, as the ERG b-wave is only an indirect measure for the photoreceptor response, we also measured the photoreceptor induced a-wave by blocking the masking ERG b-wave (Figure 4B1). This was achieved by administering a pharmacological cocktail containing the excitatory amino acid transporter inhibitor DL-threo-beta-benzyloxyaspartate (DL-TBOA) and metabotropic glutamate receptor inhibitor L-2-amino-4-phosphonobutyric acid (L-AP4) (Wong *et al*, 2004). Consistently, the double flash paradigm demonstrated that the a-wave response recovery in the evening was delayed (Figure 4B2).

In order to prove that increased response recovery times measured in the evening are a bonafide circadian event, we repeated the above experiments on larvae that were kept in constant darkness. At corresponding time points the decrease in response recovery was comparable (Figure 4C1&C2), verifying that the observed changes are regulated by an intrinsic circadian clock.

As photoresponse recovery is affected by the circadian rhythm, we hypothesized that this should also be apparent in temporal aspects of vision. Therefore, we used flicker ERG stimulation to determine the flicker fusion frequency at which individual light responses cannot be separated anymore (Figure 5A). In line with our hypothesis, we found that 9 out of 10 larvae were capable to resolve a flicker frequency of 15 Hz in the morning (Figure 5B), whereas in the evening only one out of nine larvae could resolve even a lower frequency of 10 hz. This clearly indicates that also the visual temporal resolution is under circadian control.

**Figure 5.**
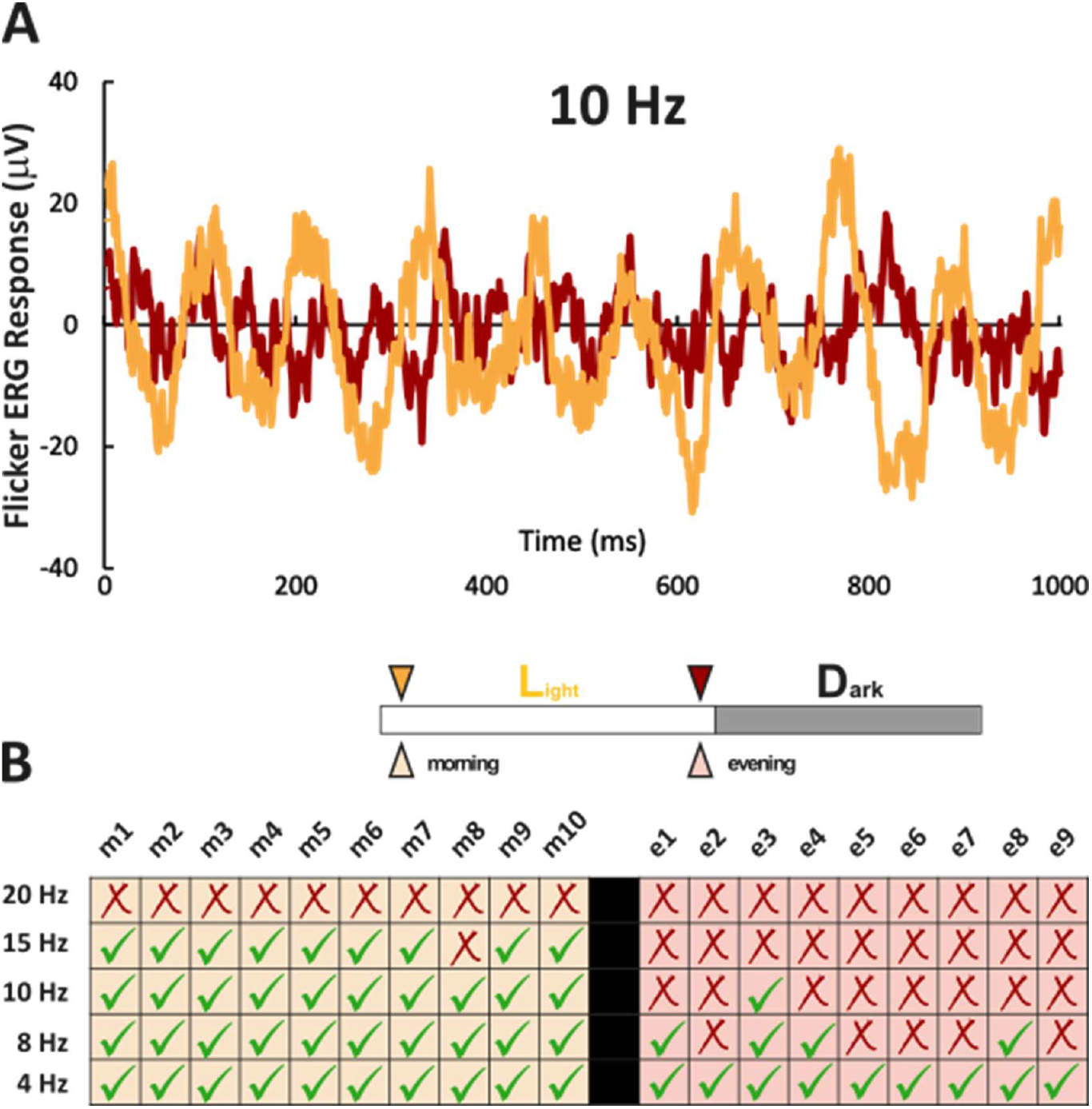
Zebrafish larvae show an increased temporal resolution in the morning. **(A)** Examples of 10 Hz flicker ERG recordings taken in the morning (yellow curve) and in the evening (red curve). Note that in the morning a temporal resolution of 10 Hz can be clearly achieved, whereas in the evening a flicker ERG of 10 Hz cannot be resolved. **(B)** Flicker response resolution table of individual larva. While the left site (light yellow) indicates the flicker ERG resolution of 10 individual larval zebrafish in the morning, the right part (light red) displays the flicker ERG resolution in the evening. “Morning” denotes recording time between ZT1 and ZT2. “Evening” denotes recording time between ZT12.5 and ZT13.5. Note that in the evening the maximal resolution was in the range of 8 Hz with only one fish being capable of resolving a 10 Hz frequency, whereas in the morning 9 out of 10 larva were capable to resolve at least 15 Hz.

### Manipulation of gene expression by light is mirrored by functional changes

Next we measured larvae reared in a reversed light cycle (DL) where the objective night turns into a subjective day. Under this condition gene expression levels stayed in the fish’s subjective time. ISH for the genes of interest (Figure 6A) reflected this, with a stronger staining intensity in LD fish in the morning compared to DL fish at the same time in the subjective evening. Consequently, when both groups were recorded at 120 hours post fertilization, a prolonged response recovery time was obtained in the fish maintained in reversed light cycle reflecting the situation in fish kept in the normal light and recorded in the evening (Figure 6D). While the intrinsic circadian clock is maintained in the absence of light, continuous light exposure has been shown to disrupt this intrinsic rhythm (Laranjeiro & Whitmore, 2014). We therefore evaluated if the circadian regulation of mRNA expression persists in larvae kept under constant light (LL). Strikingly, the gene expression differences between morning and evening detected under LD conditions were completely lost in LL larvae (Figure 6B&C). This was also reflected on a functional level with no delay of photoresponse recovery in the evening as measure by ERG.

**Figure 6.**
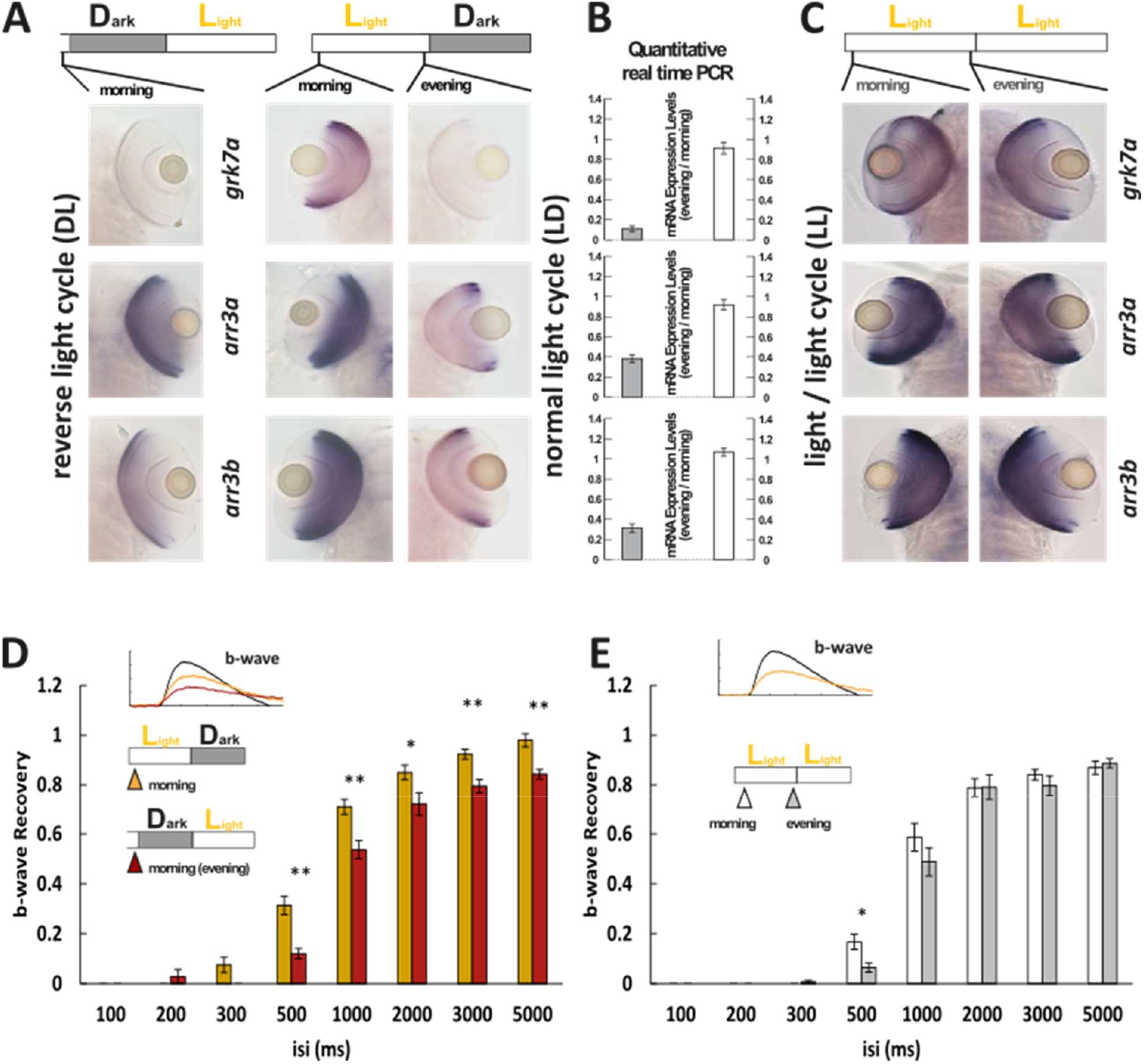
Light cycle alterations are reflected in adaptations of cone photoresponse recovery. **(A & C)** *In situ* hybridization images using arr3a, arr3b and grk7a as probes. Tissues were collected from either DL (A, left panel), LD (A, right panel) or LL (C) zebrafish larva (5 dpf) at the indicated time points. A reversal in the light cycle from LD to DL is reflected in the reversal of the *in situ* hybridization signal, with low expression levels observed in the subjective evening (A). The ratio of gene expression levels between evening (ZT13) and morning (ZT1) for fish raised under a normal light/dark (LD) cycle or under constant light (LL) are shown in (B). In contrast to the observed circadian regulation under LD conditions, under constant light (LL) conditions expression levels remain continuously elevated not displaying any circadian fluctuation (B, C). **(D)** A reversal of the light cycle is reflected in a corresponding reversal of b-wave recovery. The comparison of b-wave recovery of LD and DL larvae recorded at the same time in the morning and the subjective evening clearly indicates that immediately before darkness b-wave recovery rates are reduced. Data are presented as mean ± s.e.m (n=16 in the morning of the larvae raised in LD; n=9 in subject evening of the larvae raised in DL) of three independent experiments. Student’s t-test was used. p=0.001 at 500 ms. p=0.002 at 1000 ms isi. p=0.022 at 2000 ms isi. p=0.0009 at 3000 ms isi. p=0.002 at 5000 ms isi. * p<0.05 **p<0.01 **(E)** Under constant light (LL) conditions no changes in b-wave recovery between the subjective morning and evening can be observed. Data are presented as mean ± s.e.m (n=15 in the morning; n=12 in the evening) of three independent experiments. Student’s t-test was used. p=0.011 at 500 ms isi. * p<0.05 **p<0.01

Taken together these results demonstrate that changes in the light cycle are reflected in changes of transcript levels of phototransduction regulators that subsequently lead to altered visual performance at different times during the day.

### Visual motor response (VMR) shows difference between morning and evening

We next asked whether the observed ERG adaptations between morning and evening are directly influencing visual behavior. Therefore, we measured the visual motor response (VMR). VMR is the startle response of zebrafish larvae following a drastic change in illumination (Emran *et al*, 2008). The same individual larvae were measured in the morning (ZT1) and evening (ZT13). While the baseline activity as well as the normalized startle responses following light off is generally larger in the morning, the normalized startle responses following light-on was slightly larger in the evening (Figures 7). This suggests that the alert reaction triggered by light may be more intense when the general activity is low.

**Figure 7.**
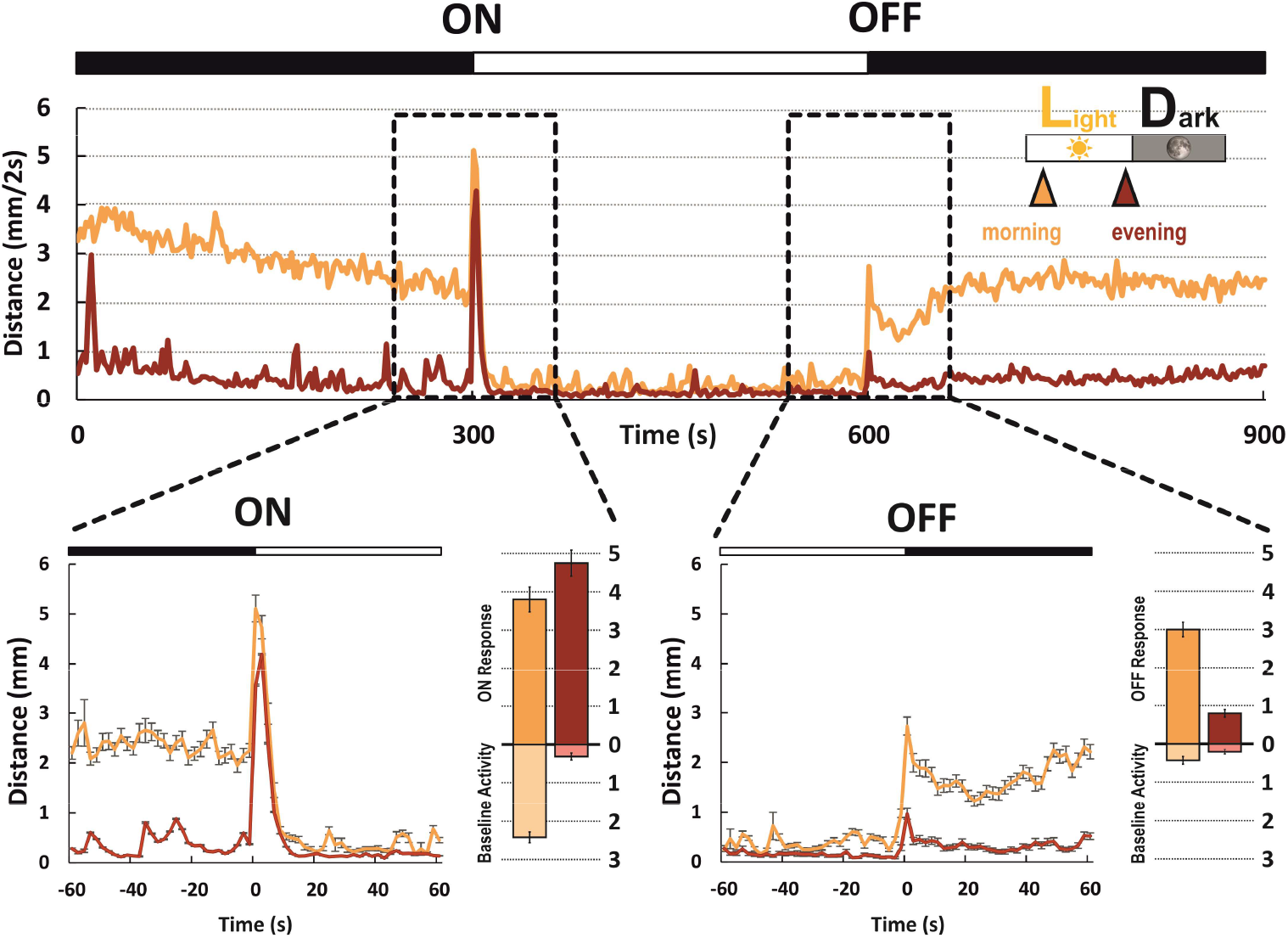
Visual behavior shows differences between morning and evening. Background movement and movement induced to sudden changes in illumination were recorded in the morning (yellow curve) and evening (red curve). Data is shown as an average of traces for fish recorded in the morning (n=190; yellow curve) and recorded in the evening (n=187; red curve) during a 15 minute period. Visual motor response (VMR) around the time of illumination changes (black dotted boxes) are enlarged. Baseline activity was quantified as the average movement in 60 seconds before the induced light change (light yellow and light red bars). Student’s t-test was used to compare the response in the morning and in the evening. Before the light-on and light off, both the baseline activities are significantly higher in the morning than in the evening, p<0.001. The amplitude of the VMR to light-on and light-off is given by the dark yellow and dark red bars. While the light-on response from fish recorded in the morning and evening have a comparable amplitude p=0.02, the amplitude of the light-off response in fish recorded in the evening is significantly reduced, p<0.001. “Morning” denotes recording time around ZT1. “Evening” denotes recording time around ZT13.

In addition to VMR, we also found that the optokinetic response (OKR) showed daily fluctuations (Figure 7-, figure supplement 1), confirming that circadian controlled expression changes in visual transduction regulators have indeed an impact on behavioral output.

## Discussion

Circadian rhythms have been shown to regulate many biological aspects of vision. An early study demonstrated that zebrafish visual sensitivity is lower before light on and higher prior to light off (Li & Dowling, 1998). Later, another study linked the rhythmic expression of long-wavelength cone opsin to the core clock component CLOCK (Li *et al*, 2008). A particularly striking finding showed that synaptic ribbons of larval zebrafish photoreceptors disassemble at night. This peculiar phenomenon may save energy in fast growing larvae (Emran *et al*., 2010). Our study now demonstrates that regulators of photoresponse decay are not only influenced by the circadian clock but in addition have a clear effect on the varying visual performances throughout a 24 hour cycle. Moreover, kinetics of cone visual transduction quenching is under the control of the circadian clock, which allows the fish to see with better temporal resolution in the morning than in the evening.

It is commonly assumed that circadian gene regulation helps the organism to optimally adapt to its preferential lifestyle and/or environment. Therefore one would expect that the circadian systems of diurnal and nocturnal animals adapt differently. Our study indeed demonstrates that orthologous zebrafish and mouse genes involved in regulating cone visual transduction decay display an anti-phasic circadian expression pattern, supporting the functional relevance of the oscillating gene expression. While the visual temporal resolution of diurnal species is reduced in the evening, the visual system of nocturnal species is tuned to be most effective during these hours. Zebrafish therefore is an interesting model to study the physiology of circadian rhythms of diurnal animals, such as humans.

We would like to point out several additional interesting observations. Although many ohnologs (paralogs generated in a whole genome duplication event) such as *grk7a* and *grk7b* share a similar circadian phase or oscillatory amplitude, others such as *rcv1a* and *rcv1b* show an almost anti-phasic relationship. This is remarkable, since these ohnologs have been generated by a teleost specific whole genome duplication event (Glasauer & Neuhauss, 2014), implying that initially all ohnologs should have been in synchronicity. Interestingly, these ohnologs also adapted different expression profiles, with *rcv1a* being expressed in rods and UV cones, while *rcv1b* is expressed in all cone types in the adult retina (UV, blue, red and green) (Zang *et al*., 2015).

While the circadian rhythmicity of most genes persists throughout all developmental stages, some genes do show markedly different expression profiles between larval and adult stages. This may be related to the fact that the larval retina is functionally cone dominant, while the adult retina is a duplex retina with rod and cone contribution. In the case of *rcv2* ohnologs, *rcv2b* displays an in-phase cyclic expression pattern throughout all stages. Conversely, *rcv2a* did not show an overt cyclic expression pattern at larval stages, but being clearly under circadian control at adult stages (Figure 1). In contrast to the *rcv1* ohnologs, both *rcv2* genes are expressed in all cone subtypes and depletion of either one acts to speed up the photoresponse termination (Zang et al., 2015). Other examples of ohnolog specific cycling have been found for *arrs* and *rgs* genes (Figure 1, Figure 1-figure supplement 4). These observations strongly indicate that the transcription of clock controlled genes (CCGs) is not uniformly regulated.

This seems hardly surprising, as around 17% of all genes expressed in zebrafish are circadian oscillating (Li *et al*, 2013). Nevertheless, only a limited number of transcription factors and corresponding binding sites which regulate the core circadian machinery have been identified so far. Although we identified some conserved binding sites of core clock proteins in our analyzed genes, neither of them was conserved in mammalian genomes (Supplementary Information & Figure 2-figure supplement 1), suggesting that the regulatory pattern of circadian regulation is more complex.

Interestingly, it has been previously demonstrated that the circadian clock seems to be desynchronized in larvae raised in darkness (Dekens & Whitmore, 2008; Kaneko & Cahill, 2005; Kazimi & Cahill, 1999; Lahiri *et al*, 2014). The circadian expression of some core clock genes and melatonin rhythms are lost when whole larvae were used as the experimental material in the absence of environmental entrainments. We did not observe this phenomenon in our study of visual transduction genes in the retina, suggesting the existence of an inheritable maternal clock in the eye (Delaunay *et al*, 2000). All the analyzed genes in our study are also expressed in the photoreceptors of the pineal gland, but the transcript fluctuations may not necessarily be synchronized between eye and pineal gland (Figure 1-figure supplement 1). Among the studied genes in zebrafish, *grk7a* expression level increased by around 50 times in one day (Figure 1A), whereas Grk7a protein only raised by about 2 times in a 24-hour-period (Figure 3A). *arr3a* transcript increased about 10 times (Figure 1C), while its protein level only grew less than 50% throughout the day (Figure 3A). Therefore these mRNA expression levels reflect proportionally to protein levels, indicative of a rather fast turnover rate for these proteins.

Our OKR experiments revealed that in the evening eye velocity was reduced when compared to measurements at noon. However, between morning and evening identical results were obtained, despite slower photoresponse recovery in the evening (Figure 5-figure supplement 1). This was also true for larvae raised in a reversed light cycle and measured at an identical stage, indicating that maturation cannot be the cause of the observed results. Another explanation is that visual transduction quenching rate may peak at another time rather than the time of our recordings. Since the OKR is a very robust response, the difference in the visual transduction may be too subtle to produce a visible change in OKR between morning and evening.

In conclusion, we have shown that key regulators of cone visual transduction at both mRNA and protein level are under circadian control. Moreover, expression levels of these regulators in diurnal and nocturnal species are anti-phasic, suggesting that circadian changes influencing physiological and behavioral properties of vision are reflected in adaptation to different visual ecologies.

## Materials and Methods

### Zebrafish care

Usually zebrafish (*Danio rerio*) were maintained at a standard 14 h light: 10 h dark cycle (LD) with light on at 8 am and light off at 10 pm. DL (light on at 8 pm and light off at 10 am), DD (constant darkness; starting at 2 hpf) and LL (constant light) fish were raised in separate incubators. Water temperatures were always kept between 26 and 28 °C (Amores *et al*, 1998). Only fish from the WIK wildtype strain were used in our study. Embryos were raised in E3 medium (5 mM NaCl, 0.17 mM KCl, 0.33 mM CaCl2, and 0.33 mM MgSO4) containing either 0.01% methylene blue to suppress fungal growth and/or 0.2 mM PTU (1-phenyl-2-thiourea; Sigma-Aldrich) to prevent pigment development. Adult zebrafish were sacrificed using ice water following decapitation. All animal experiments were carried out in the line with the ARVO Statement for the Use of Animals in Ophthalmic and Vision Research and were approved by the Veterinary Authorities of Kanton Zurich, Switzerland (TV4206).

### Zebrafish Quantitative Real-Time PCR (qRT PCR)

Around 30 5 dpf larvae or 5 eyeballs from adult zebrafish were collected per time point (ZT 1, 4,7,10,13,16,19 and 22) and the tissue stored in RNAlater (Sigma) at 4°C. Dark adapted tissue was collected under dim red light. Only eyeballs were used for RNA extraction using the NucleoSpin® RNA kit (Macherey-Nagel). cDNA was produced using 110 ng total RNA as template for reverse transcription with SuperScript® III (Invitrogen, Life Technologies, Zug, Switzerland). qRT-PCR (Applied Biosystems Prism SDS 7900HT; Life Technologies) was performed using the MESA Green qPCR Mastermix Plus for SYBR Assay (Eurogentec,Seraing, Belgium) on a liquid handling robot platform (Tecan Genesis). Primers (Sigma-Aldrich) for qRT-PCR were intron-spanning to avoid amplification of non-digested genomic DNA fragments and were designed by online Universal ProbeLibrary Assay Design Center (Roche). Standard housekeeping genes (elongation factor 1, ef1; β-actin2, actb2; and ribosomal protein L 13, rpl13) were used as reference (Tang *et al*, 2007). Expression levels were normalized to 1.

### Mouse care and gene expression analysis

Mice were maintained at the Laboratory Animal Services Center (LASC) of the University of Zurich in a 12 h light: 12 h dark cycle with lights on at 7 am. All animal experiments were performed according to the ARVO Statement for the Use of Animals in Ophthalmic and Vision Research and the regulations of Veterinary Authorities of Kanton Zurich, Switzerland.

10 to 12-week-old wildtype mice (129S6, Taconic, Ejby, Denmark) were used in our experiments. Dark phase mice were killed under red light and retinas were processed further under normal light conditions. 3 mice at each time point (ZT 1, 5, 9, 13, 17, 21) were sacrificed and RNA was extracted (Macherey-Nagel, Oensingen, Switzerland) according to manufacturer’s instruction. cDNA synthesized using oligo-dT was done as previously described (Storti *et al*, 2019). qRT-PCR was performed by ABI QuantStudio3 machine (Thermo Fisher Scientific) with the PowerUp Sybr Green master mix (Thermo Fisher Scientific). Primer pairs used are listed in Table2 for each gene of interest. Beta-actin (*Actb*) was used as a housekeeping gene to normalize gene expression with the comparative threshold cycle method (DDCt) using the Relative Quantification software (Thermo Fisher Scientific). The highest expression level was normalized to 1.

### *In Situ* Hybridization (ISH)

Primers used to generate *in situ* probes are listed in table 3. Probes were digoxigenin-labeled using the DIG RNA Labeling Mix purchased from Roche.

**Table 3.**
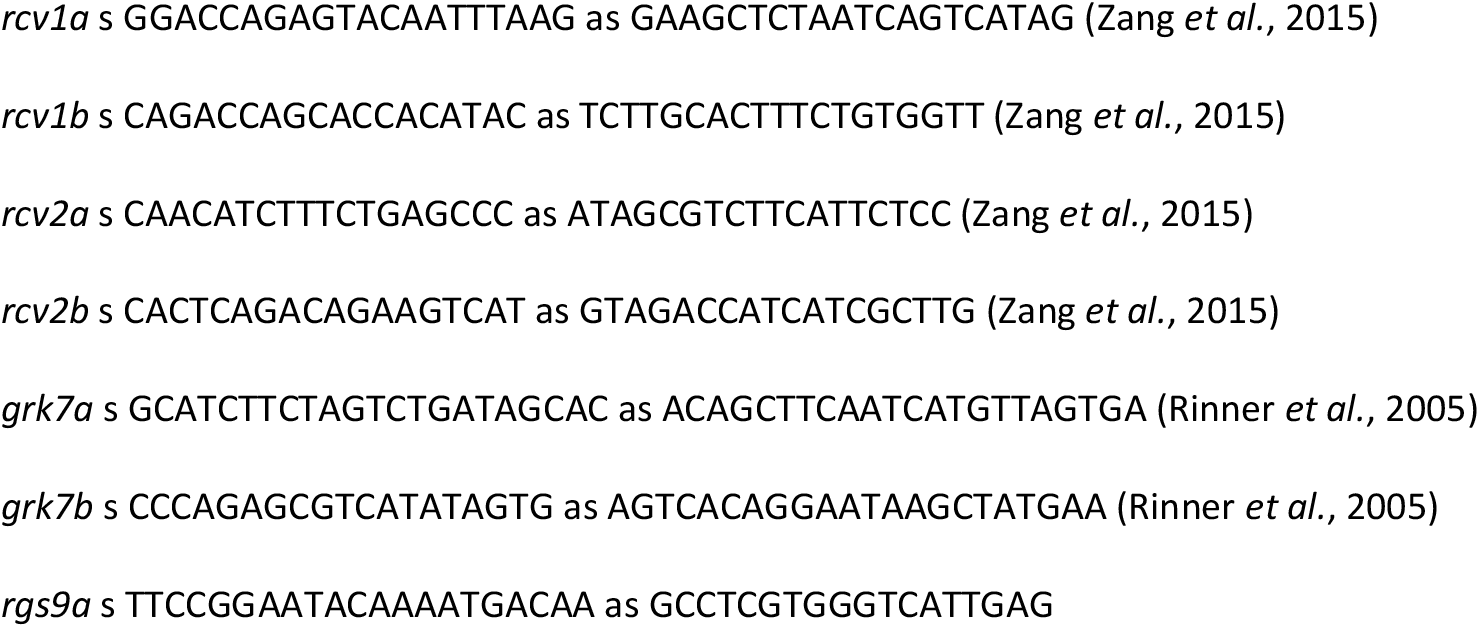
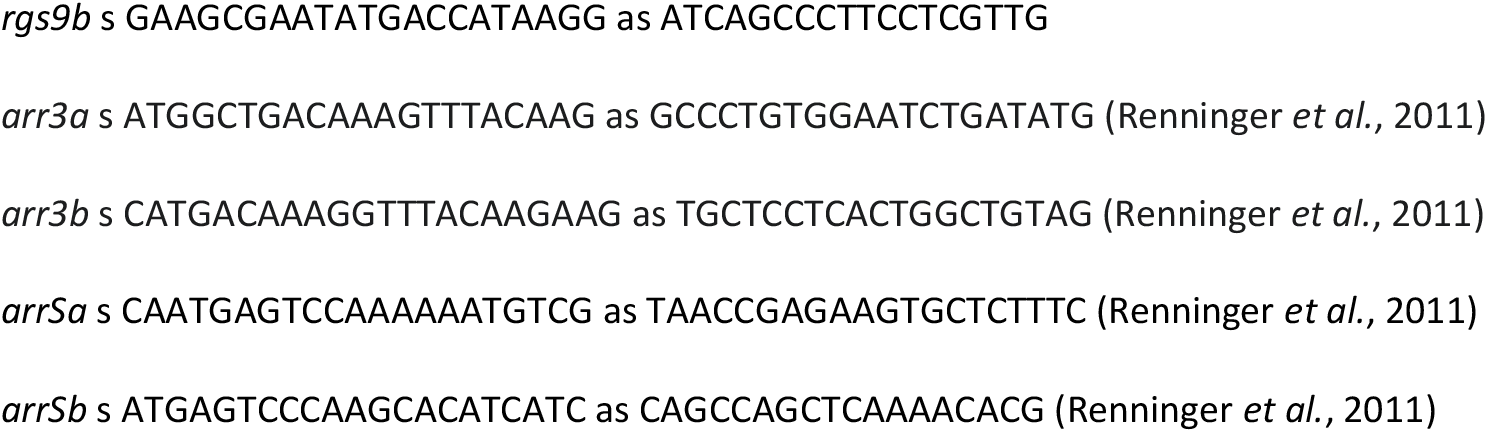
Primer sequences for *in situ* probe preparation.

For whole-mount ISH embryos were treated with E3 containing 0.2mMPTU (1-phenyl-2 thiourea; Sigma-Aldrich) to avoid pigmentation. 5 dpf larvae were fixed in 4% paraformaldehyde (PFA; Sigma) in phosphate-buffered saline (PBS) overnight at 4 °C. Time points with maximal differences were chosen according to qRT-PCR results. Embryos were washed 3 times in PBS containing 1% Tween (PBT), dehydrated step-wise (25%, 50%, 70% MeOH in PBT) and stored in 100% MeOH at -20 °C.

For slide ISH, eyeballs were removed from adult zebrafish at different time points. And fixed overnight at 4 °C using 4% PFA. Detailed ISH processes have been previously described (Haug *et al*, 2015).

### Infrared Western Blotting

5 eyeballs from adult zebrafish were homogenized in ice-cold 150 ml RIPA buffer (150 mMNaCl, 1%Triton-X, 0.5% sodiumdeoxycholate, 50 mM Tris (pH 8), 1 mM EDTA, 0.1% SDS) containing cOmplete™ Protease Inhibitor Cocktail (Roche). After 2 h incubation on a 4°C shaker, lysates were centrifuged for 30 min at 4°C. Supernatants were stored at -80°C. Nitrocellulose membranes with 0.45 µm pore size were used. Primary antibodies were diluted to the following concentrations: rabbit anti-Arr3a: 1 : 4,000; rabbit anti-Grk7a: 1 : 3000; mouse anti-β-Actin: 1 : 6000 (Renninger *et al*., 2011; Rinner *et al*., 2005). Anti-arr3a and anti-β-Actin antibody or anti-Grk7a and anti-β-Actin antibody were applied simultaneously. Secondary antibody IRDye® 800CW Goat anti-Rabbit IgG and IRDye® 680RD Goat anti-Mouse IgG (LI-COR) were diluted 1:20,000 in blocking buffer (1% BSA in PBST). Signal was detected by the Odyssey® CLx Imaging System (LI-COR) and the data was normalized to the internal loading control β-Actin by IMAGEJ (Schindelin *et al*, 2012).

### Electroretinography (ERG)

ERG was recorded as previously described (Zang *et al*., 2015). Light intensity (light source: Zeiss XBO 75 W) was measured using a spectrometer (Ocean Optics, USB2000b; software Spectra Suite, Ocean Optics) with a spectral range describe previously (Zang *et al*., 2015). Pairs of two light flashes with equal intensity and duration (500 ms) were applied (Rinner *et al*., 2005). Intervals between two flashes were either 100, 200, 300, 500, 1000, 2000, 3000 or 5000 ms. The interval between two pairs was 20 s. b-wave recovery is defined as the ratio of the second b-wave amplitude to the first one in the same pair.

To measure ERG a-wave, 5 dpf larval eyeballs were treated with 400 µM APB and 200 µM TBOA in Ringer’s solution (111 mM NaCl, 2.5 mM KCl, 1 mM CaCl2, 1.6 mM MgCl2, 10 μm EDTA as chelator for heavy metal ions, 10 mM glucose, and 3 mM HEPES buffer, adjusted to pH 7.7–7.8 with NaOH). A HPX-2000 (Ocean Optics) light source was used. Intervals between two flashes were 300 ms, 500 ms, 1000ms and 1500 ms respectively. a-wave recovery is defined as the ratio of the second a-wave amplitude to the first one in the same pair.

Flicker fusion ERGs were measured with a white light LED bulb that was connected to a pulse generator (Grass). Each pulse lasted for 15 ms and the flicker frequencies of 4 Hz, 5 Hz, 8 Hz, 10Hz, 15 Hz and 20 Hz were used. Each flicking light stimuli was presented for 5 s.

### Visual motor response (VMR)

The visual motor response (VMR) was measured using a Zebrabox (ViewPoint Life Science, Lyon, France). 5 dpf larvae were placed in a 96-well plate, dark adaptation for 10 minutes inside the Zebrabox and larval movement recorded with light off, on and off for 5 minutes each. The distance which a single larva moved was measured every 2 seconds. Baseline activity was calculated as the average of movement 1 minute before light on or off.

### Optokinetic response (OKR)

The OKR was recorded as previously described (Rinner *et al*., 2005). Briefly, 5 dpf larvae were tested with sinusoidal gratings at different time points (ZT 1, 4, 7, 10 and 13). To determine the contrast sensitivity a spatial frequency of 20 cycles/360° and an angular velocity of 7.5 deg/s were used with different contrast settings (5%, 10%, 20%, 40%, 70% and 100%). To explore the spatial sensitivity, an angular velocity of 7.5/s and 70% of maximum contrast were applied with varying spatial frequency (7, 14, 21, 28, 42, and 56 cycles/360°). Figures were prepared by SPSS (Version 23.0. Armonk, NY: IBM Corp).

## Acknowledgements

We like to thank Kara Kristiansen and Martin Walther for expert animal maintenance.

---

**Figure 1-figure supplement 1.**
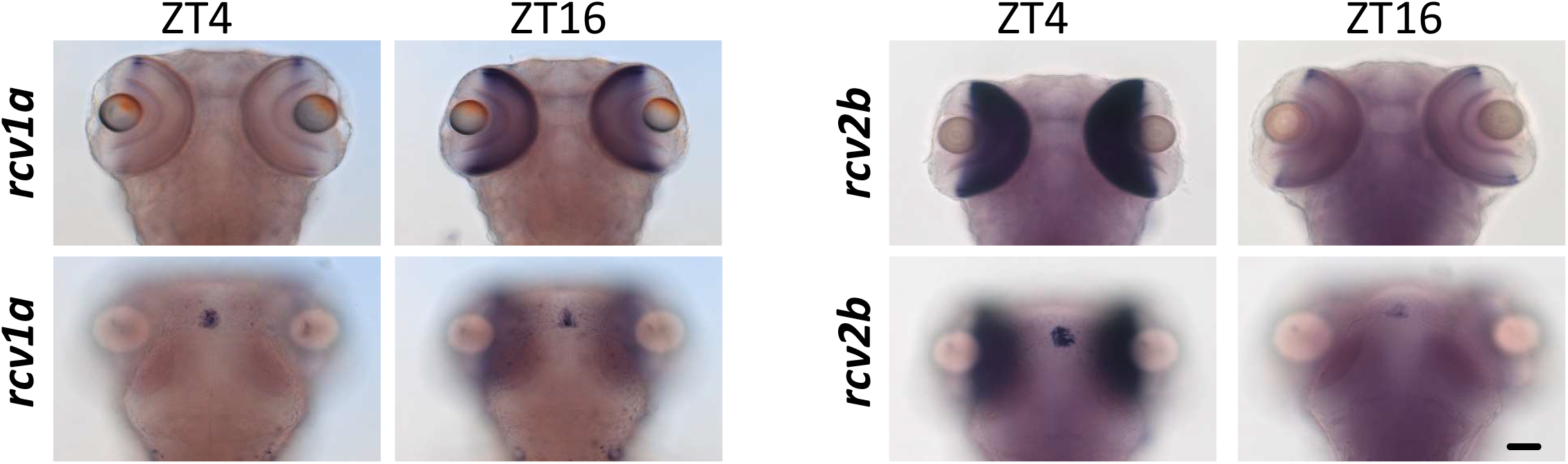
ISH of rcv1a and rcv2b as examples indicating the staining in pineal gland may not or may be synchronized with the staining in the eye. Scale bar (=50 µm) applies to all panels.

**Figure 1-figure supplement 2.**
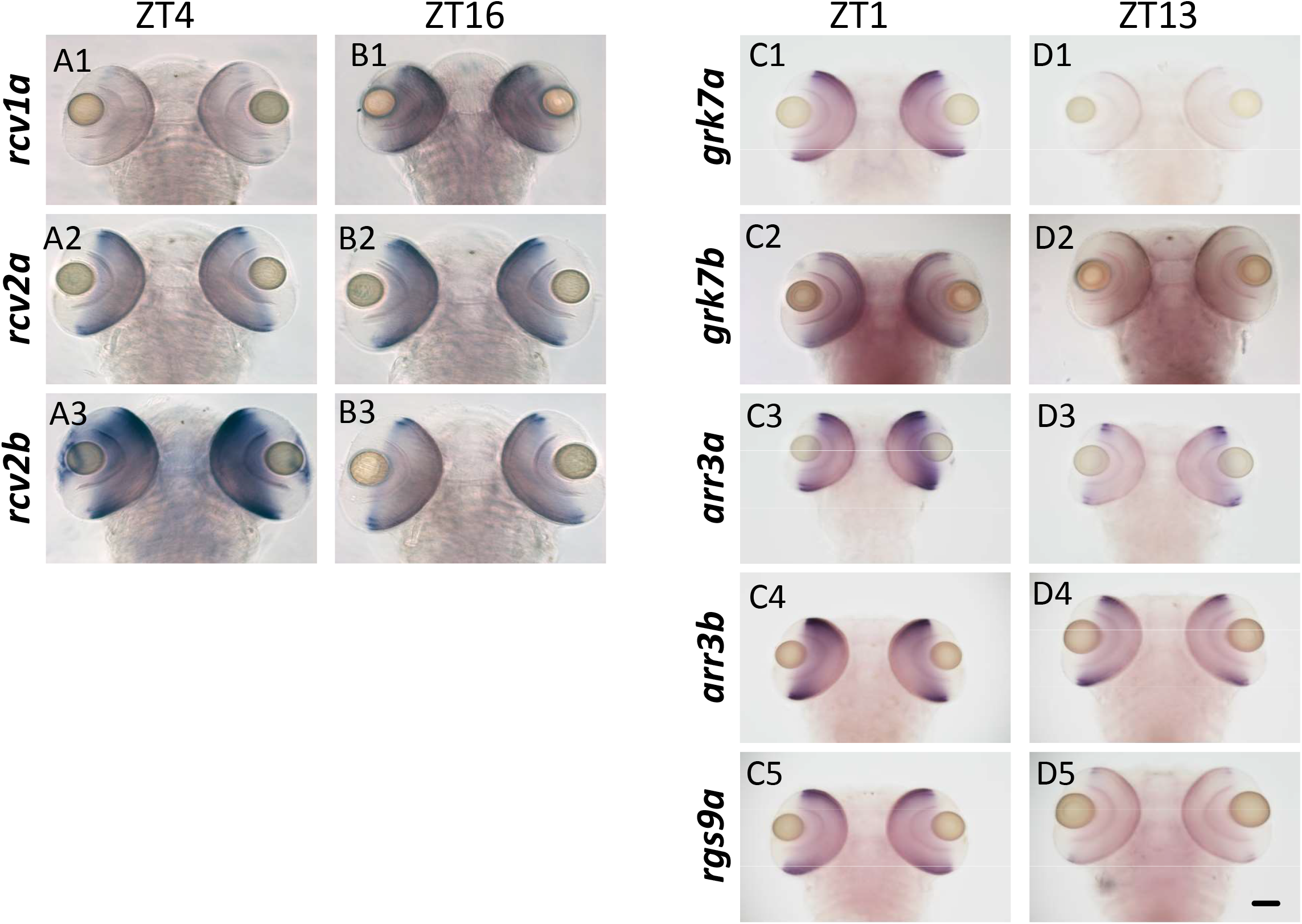
ISH showing different gene expressions at varying time points in 5 dpf zebrafish larvae in dorsal view. Scale bar (=50 µm) applies to all panels.

**Figure 1-figure supplement 3.**
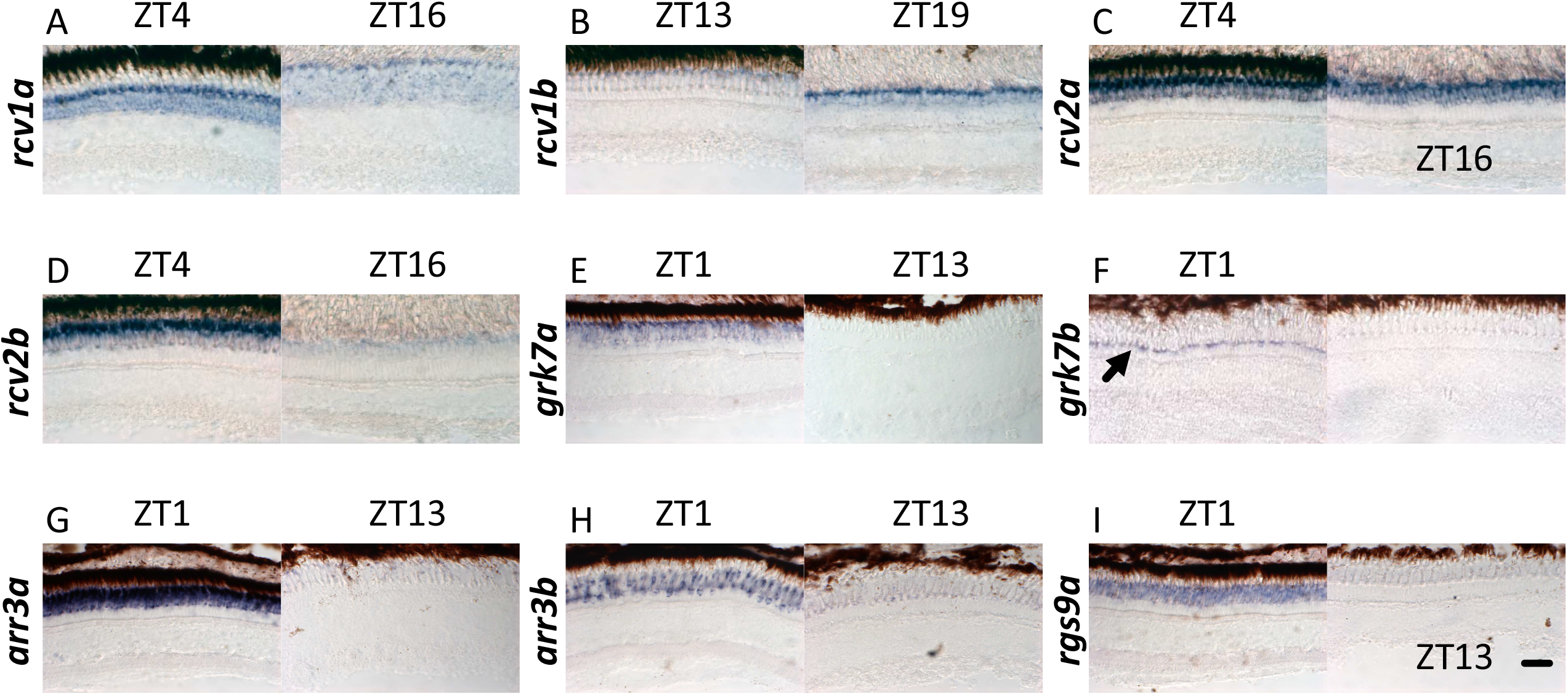
ISH showing different gene expressions on radial sections of adult zebrafish retina at different time points indicated on top. Arrow denotes UV cones. Scale bar (=20 µm)

**Figure 1-figure supplement 4.**
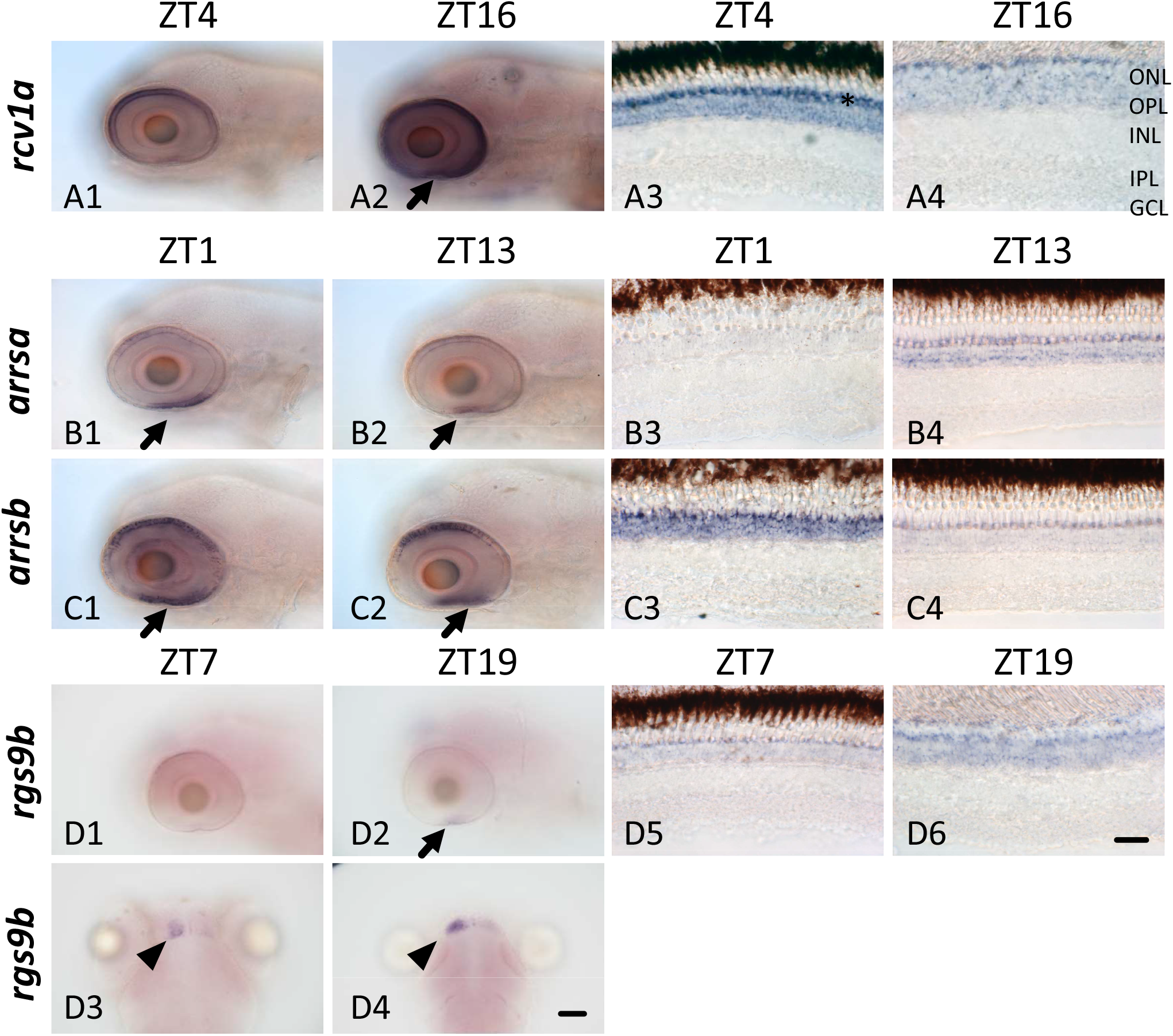
ISH showing different rod gene expressions in zebrafish larvae and adult retina at different time points indicated on top. Arrows denote region enriched with rods. Arrowhead denotes habenula. Star denotes UV cone layer. GCL, ganglion cell layer; INL, inner nuclear layer; IPL, inner plexiform layer; ONL, outer nuclear layer; OPL, outer plexiform layer. Scale bar (=20 µm) applies to corresponding panels.

**Figure 2-figure supplement 1.**
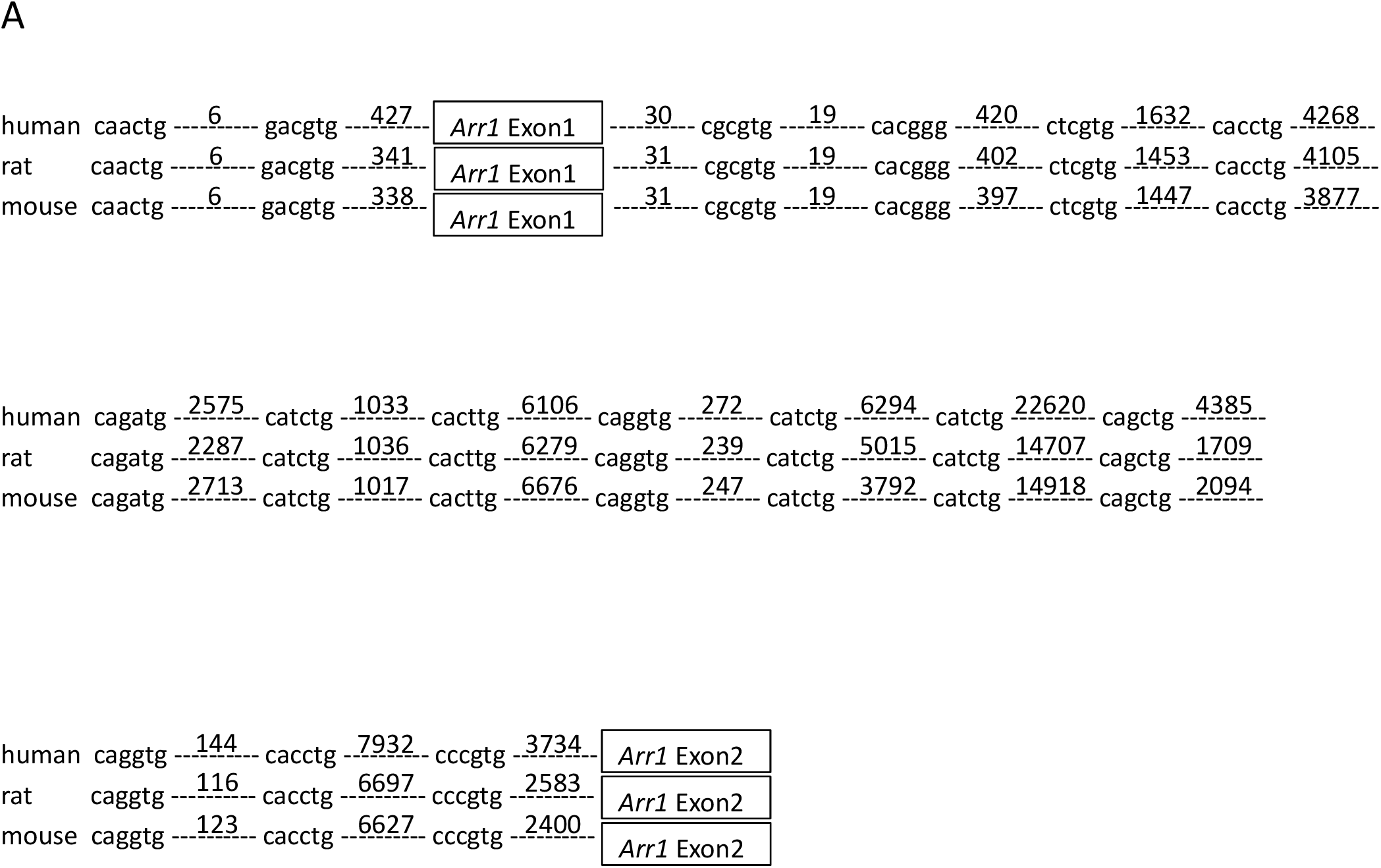
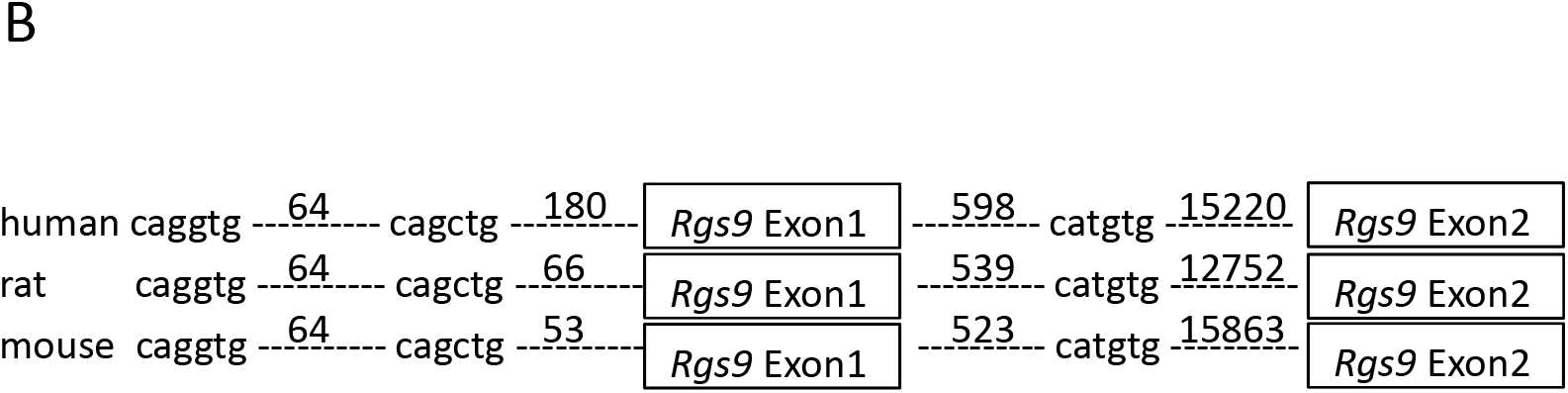
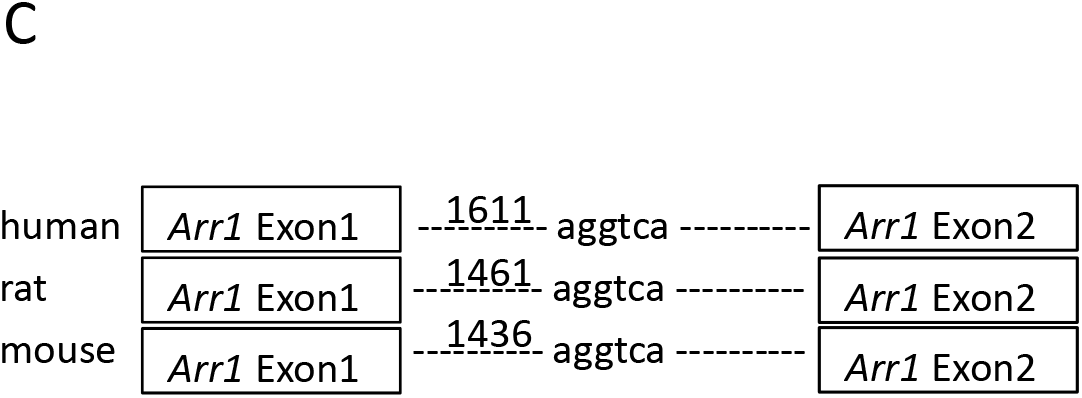
Conserved non-canonical E-boxes in mouse genes.

**Figure 7-figure supplement 1.**
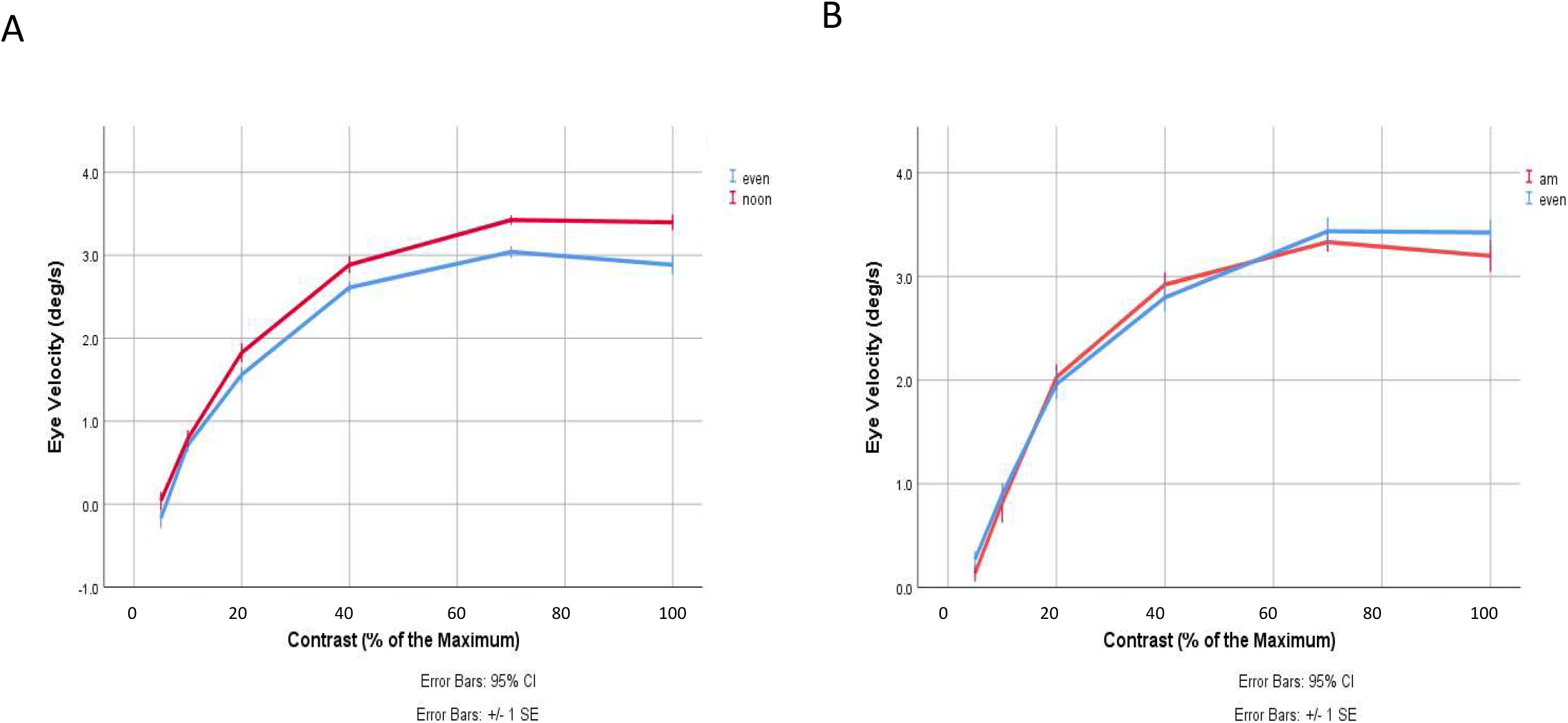
(A)OKR measurements indicate increased contrast sensitivity at noon compared to evening but similar contrast sensitivity between morning and evening. Eye velocity as a function of stimuli contrast recorded at noon and in the evening (repeated measures ANOVA by SPSS (IBM, version 26.0) p=0.001). “Noon” denotes recording time around ZT5, n=14. “Evening” denotes recording time around ZT13, n=21. Data are presented as mean ± standard deviation (SD) of three independent experiments. (B) Eye velocity as a function of stimuli contrast recorded in the morning and in the evening. (Repeated measures ANOVA by SPSS (IBM, version 26.0) p=0.671). “am” denotes recording time around ZT1, n=12. “Evening” denotes recording time around ZT13, n=10. Data are presented as mean ± SD of three independent experiments

